# Translational control of *furina* by an RNA regulon is essential for heart morphogenesis and cardiac valve function

**DOI:** 10.1101/2023.01.30.526270

**Authors:** Agnieszka Nagorska, Finnlay Lambert, Angus Inman, Sara Toral-Perez, Andreas Zaucker, Jan Gorodkin, Wan Yue, Michael Smutny, Karuna Sampath

## Abstract

Heart development is a complex process, starting from specification of cardiac precursors and formation of a linear tube to gradual progression to a functional beating organ. For normal heart development, many processes, including asymmetric positioning of the heart along the left-right (L/R) axis, cardiac growth, and cardiac valve morphogenesis must be completed successfully. Although heart development has been studied extensively, the mechanisms that control heart morphogenesis and valve formation are not fully understood. The pro-convertase FurinA is a key protein that functions in heart development in many vertebrates including zebrafish. How FurinA activity is regulated during heart development is not known. Through computational analysis of the zebrafish transcriptome, we identified a short sequence and structure RNA motif in a variant transcript of FurinA harbouring a long 3’untranslated region (3’UTR). The alternative 3’UTR *furina* isoform is expressed at embryonic stages preceding organ positioning. Reporter localization and RNA-binding assays show that the *furina* 3’UTR forms complexes with the conserved RNA-binding protein and translational repressor Ybx1. Conditional mutant zebrafish embryos affecting *ybx1* show premature and increased Furin reporter protein expression, abnormal cardiac morphogenesis and heart looping defects. Our mutant *ybx1* hearts have an expanded atrioventricular canal, abnormal sino-atrial valves and many mutant embryos show retrograde blood flow from the ventricle to the atrium. This is similar to human heart valve regurgitation patients. Our findings show an essential function for the 3’UTR element/Ybx1 regulon in translational repression of FurinA, revealing a new upstream regulatory mechanism that controls embryonic heart development, and demonstrates the *ybx1* mutant as a model to study cardiac valve development and function.

## Introduction

Cardiovascular disease is a leading cause of death and accounts for a quarter of all deaths in the UK (Cheema *et al*., 2022). Even though cardiovascular development has been extensively studied, the molecular mechanisms underlying congenital heart disease are not fully known. The importance of RNA-binding proteins (RBP) and RNA motifs in regulating organ development and diseases has been recently uncovered and understanding their interactions is fundamental to developing future diagnostic and therapeutic strategies.

Zebrafish is a popular research model for studying heart development because the embryos are transparent, allowing in-depth observation of formation of many internal organs using live microscopy until larval stages. Additionally, although the heart in zebrafish embryos develops by 24 hours post-fertilisation (hpf), embryos can survive the first week of life without a circulatory system. Zebrafish heart development is a dynamic process: at 24 hpf, a primitive beating organ is already formed and by 48 hpf, looping of the chambers and atrioventricular (AV) canal formation is complete. Although heart formation occurs rapidly, many complex processes must be completed for the functional organ to develop. One key process is the establishment of the left-right asymmetry, which is required for the correct placement of internal organs including the heart (Bakkers, 2011).

In mammals, positioning of the heart on the left side and liver on the right is referred to as *situs solitus*. Disruptions to this process may cause left or right isomerism disorders that affect ∼1:10,000 individuals (Blum and Ott, 2018). Following heart looping, morphogenesis of the valves is initiated in the atrioventricular canal by formation of the endocardial cushions that gradually differentiate and form functional heart valves (Gunawan *et al*., 2019). At similar stages, zebrafish embryos are small enough to obtain the required oxygen to live by passive diffusion. Therefore, unlike other model organisms, many mutations in the zebrafish heart do not cause embryonic lethality within the first week of zebrafish life (Liu and Stainier, 2012). This is especially useful for studying key processes in cardiac differentiation, and cardiac valve and septal development, which if disrupted in mammals result in foetal death.

The mechanisms that control left-right asymmetry are well-conserved in vertebrates. For instance, asymmetric left-sided expression of the secreted growth factor Nodal during somitogenesis in the lateral plate mesoderm (LPM) has been reported in many organisms including frogs, fish, chick, and mice (Levin *et al*., 1995; Lowe *et al*., 1996; Sampath *et al*., 1997, 1998; Varlet and Robertson, 1997; Long, Ahmad and Rebagliati, 2003; Shiratori and Hamada, 2006). Motile cilia generate calcium transients in the left-right organiser, and drive leftward Nodal gene expression (Francescatto *et al*., 2010; Shinohara and Hamada, 2017). Expression of the Nodal inhibitor Lefty1 in the embryonic midline spatially restricts Nodal to the left LPM and results in asymmetric expression of the Nodal target, Pitx2 (Lowe *et al*., 1996; Lin *et al*., 1999; Thisse and Thisse, 1999; Chen and Shen, 2004; Shiratori and Hamada, 2006; Schottenfeld, Sullivan-Brown and Burdine, 2007).

The pro-protein convertase FurinA cleaves Nodal and is essential for heart development in many vertebrates (Constam and Robertson, 2000; Beck *et al*., 2002). Studies in zebrafish showed that FurinA is essential for maturation of the Nodal homolog Southpaw (Spaw) and establishes its signalling range in the LPM. Mutations disrupting zebrafish FurinA result in heart looping and trabeculation defects. The *furina* mutant embryos also display abnormal jaw development and 80% of maternal zygotic mutants die due to lack of inflation of the swim bladder (Walker *et al*., 2006; Tessadori *et al*., 2015; Zhou *et al*., 2021). In the mouse, deletions in Furin cause embryonic lethality, and the mutants display a range of cardiac morphogenesis defects including ventral closure, abnormalities of outflow tract and severe heart looping defects. Furin mutant mice also exhibit impairment of large vessel formation and defects in yolk sac vasculature (Roebroek *et al*., 1998; Dupays *et al*., 2019). Conditional deletion of *Furin* in endothelial cells in mouse did not reduce the severity of phenotype, and heart valve malformations in this allele also resulted in embryonic lethality (WooJin *et al*., 2012), underscoring the importance of Furin during cardiac development.

Although the molecular mechanisms underlying transcriptional control of the Nodal pathway have been extensively studied, very few studies have explored regulation of the left-right axis by non-coding regions and post-transcriptional regulation (Minegishi *et al*., 2021).We found that many Nodal pathway components in zebrafish have a short, conserved 3’UTR motif characterised by a AGCAC sequence motif, followed by a stem and a loop. The 3’UTR element was first identified in *squint/nodal-related 1(sqt/ndr1)* transcripts. A fluorescent *sqt* RNA reporter localises to dorsal progenitors in early zebrafish embryos (Gore *et al*., 2005; Gilligan *et al*., 2011; Kumari *et al*., 2013). We demonstrated that transcripts of the Nodal inhibitors, *lefty1* and *lefty2*, also contain the 3’UTR motif, and lefty reporter RNAs also localize in early embryos, indicating similar activities of the 3’UTR motifs (Zaucker *et al*., 2017). The conserved RNA-binding protein Ybx1, binds the 3’UTR element in *sqt, lefty1* and *lefty2* and translationally represses these RNAs during early development (Kumari *et al*., 2013; Zaucker *et al*., 2017). Overexpression of full length lefty1/2 GFP versus lefty1/2 lacking the 3’UTR motif showed that deletion of the element leads to a strong loss-of-nodal phenotype through inhibition of Nodal (Zaucker *et al*., 2017). Mutations in mammalian Ybx1 also led to de-repression of Nodal and Lefty in cultured cells, suggesting broader conservation of this mechanism.

Since Ybx1 was found to translationally repress a range of different mRNAs through the 3’UTR motif, the motif was named ‘Ybx1 binding element’ (YBE) (Zaucker *et al*., 2019). By computational analysis of the zebrafish transcriptome, we now find that YBE motif is present in a variant 3’UTR transcript (variant X1) of the Nodal maturation enzyme, FurinA, but not in *spaw/ndr3*. We show that the *furina* variant X1 harbours the YBE motif in its 3’UTR and is expressed prior to organ positioning in zebrafish. Conditional *ybx1* mutant embryos reveal left-right axis defects and abnormal visceral organ positioning. In addition, these *ybx1* mutant embryos have an enlarged AV canal and show retrograde blood flow. This is similar to mitral valve regurgitation disorder, a condition characterised by a leaky heart valve and back-flow of blood into the heart chambers of human patients. Our work shows that translational control by Ybx1 functions in left-right axis formation, heart morphogenesis and valve development, and identifies a new zebrafish model for mitral valve regurgitation disorders.

## Methods Zebrafish strains

Zebrafish were maintained in accordance with the University of Warwick institutional animal welfare regulations and the UK home office guidelines. All fish in this work were maintained in the Tübingen (TU) background. Zebrafish *ybx1* ^*sa42*^ fish lines and controls were set up as single pair crosses separated with a divider the previous day, and on the next morning, the divider was removed to allow mating. Embryos they were kept at 28°C in 0.3X Danieau’s solution (17 mM NaCl, 2 mM KCl, 0.12 mM MgSO_4_, 1.8 mM, Ca(NO_3_)_2_, 1.5 mM HEPES, pH 7.6) until 50% epiboly, and afterwards they were shifted to 22°C until the 20-somite stage. Embryos were then shifted back to 28°C until 50 hpf (long-pec) for live imaging or fixed in 4% PFA, dehydrated and stored in methanol at -20°C.

### PCR and qPCR amplification

Zebrafish embryos from stages 1K, 50% epiboly, 10 somites, 18 somites and adult ovaries were collected and 50 eggs/embryos were used per stage. RNA was initially homogenised in TriZol reagent (Thermofisher) and then extracted using RNA miniprep kit (NEB). Integrity of purified RNAs was checked on agarose gel and concentration was measured on the nanodrop. Next RNAs were reverse transcribed using superscript III (Thermofisher). For qRT-PCRs, each cDNA was amplified using GAPDH, furina and furina X1 primers in a standard 20 uL PCR reaction (5x go taq green buffer Promega, dNTPs 10mM Promega, of 2µm each forward and reverse primer, home-made taq polymerase and sterile water). For qPCR we used the 2x Luna Universal qPCR master-mix (NEB), following the manufacturer’s protocol. All primers were tested for efficiency to ensure they were comparable. Primer list available in supplementary data.

### In situ hybridization

Embryos were fixed at the required developmental stages overnight at 4ºC in 4% paraformaldehyde in PBS (phosphate buffered saline) and whole mount *in situ* hybridization (WISH) was performed according to established protocols (Thisse and Thisse, 2008).

### Fluorescent RNA synthesis and injections

Plasmids were linearized using Not1 restriction enzyme and mRNA was transcribed from purified template DNA using SP6 RNA polymerase. Transcription master-mix contained 500 ng of linear purified template, 5x transcription buffer, DTT, ribonuclear capping mix (rATP, rGTP, rCTP 10mM each) rUTP (100mM), 7mG(5’)pppG cap (7.5 mM), and Alexa 488-5-UTP (1mM) or Alexa 546-UTP (1mM) (Thermofisher) fluorophore. Following transcription samples were treated for 30 minutes with a Turbo DNase I enzyme and unincorporated nucleotides were removed using mini-spin columns (Bio-Rad). Next, phenol-chloroform purification was performed on the fluorescent RNAs, followed by overnight precipitation in isopropanol and elution in nuclease free water. RNA integrity was checked on an agarose gel and the concentration and incorporation rate of the fluorophore was checked using nanodrop. Embryos were injected at the 1-cell stage with 10-25 pg aliquots of fluorescent mRNAs, imaged and/or scored at 4 cell stage.

### RNA Immunoprecipitation

Zebrafish embryos were collected at following stages: 50 % epiboly, 10-somites and 18-somites in aliquots of 250 embryos. Embryos were cross-linked in 1% formaldehyde solution for 20 minutes, rinsed 3 times with PBS-T and lysed using RIPA buffer in DEPC water (50 mM Tris–Cl pH 7.5, 1% NP-40, 1% sodium deoxycholate, 0.1% SDS, 1mM EDTA, 1 M NaCl, 1 M urea, protease inhibitor). Immunoprecipitation was performed according Kumari *et al* 2013 and Zaucker *et al* 2018, with an anti-Ybx1 antibody (Sigma 4F12). A list of primers used in RT-PCRs and qPCRs is available in the supplementary material.

### Immunofluorescence

Antibody staining was performed as described in Santos *et al* 2018. The MF20, S46 and ZN-8 antibodies were purchased from Developmental Studies Hybridoma Bank and used according to the manufacturer’s recommendations.

### Translation assays and Western blot analysis

Full length *spaw*-GFP and *furina* sfGFP were cloned into pCS2 vector by Gibson assembly and transcribed using the SP6 mMachine kit (Thermofisher). RNAs were treated with turbo DNase, phenol-chloroform purified, precipitated overnight in isopropanol and eluted in nuclease free water. Next 50 pg of RNA was injected into 1 cell stage ybx1^sa42/sa42^ mutant embryos, and 50% of the injected embryos were temperature shifted to 22°C at the 16-cell stage, whereas control embryos were kept at 28°C. At the 512-cell stage all embryos were collected and lysed in RIPA buffer (50 mM Tris-Cl pH 7.5, 1% NP-40, 0.5% NaDeoxycholate, 0.05% SDS, 1mM EDTA, 150 mM NaCl) with the proteinase inhibitor (Roche). The sample concentration was normalised using Bradford assay kit (NEB), following which the samples were boiled for 5 min in Laemmli buffer, loaded on a 10% acrylamide gel and western blot was carried out. Membranes were incubated with 1 in 2000 anti-GFP HRP conjugated antibody (Bio-Rad) and 1 in 4000 HRP conjugated Actin loading control (Santa Cruz).

### Confocal imaging

Imaging of the immunofluorescence and translation assay was done using the Andor Revolution Spinning Disk system, with a Yokogawa CSU-X1 spinning disk unit, fitted with a 488 nm laser and captured with an iXon Ultra 888 EMCCD camera. Acquisition of the Z stacks for image was done with Andor software and final processing and analysis in ImageJ.

### Heart imaging

MZ*ybx1* and wild type control embryos were collected at the same time and grown at 28.5 ºC until the 75% epiboly stage. They were then temperature shifted to 22 ºC until the 21-som stage and shifted back to 28ºC until 5 dpf. At 24 hpf, embryos were treated with the phenylthiourea (PTU) at 0.003% concentration in Danieau’s solution to prevent pigment formation. Prior to imaging on day 5, embryos were immobilised by adding 25× tricaine solution to Danieau’s solution and mounted in 0.6% low melt agarose on a 3-cm glass cover slip-bottom petri dish for imaging at 13 frames per second using Nikon ECLIPSE Ni microscope with a HAMAMATSU digital camera C11440, ORCA-Flash4.OLT. Time-lapse movies of approximately 30 seconds-1 minute duration were acquired.

To analyse the direction and velocity of the blood flow in *ybx1* and control embryos, Particle image velocimetry (PIV) application was used in MATLAB (Thielicke, W. and Sonntag, 2021). Each movie was converted in ImageJ using the function ‘find edges’ and uploaded to PIV, the region of interest was set out to be a junction in the AV canal and frames were analysed. Vectors were saved and analysed for direction with a custom MATLAB code.

### RNA Structure probing

In vitro probing of RNAs were probed as described in (Flynn *et al*., 2016), except that MgCl2 concentration for 3.3× SHAPE reaction buffer was reduced to 3.3 mM and probing reactions were conducted at 28 °C. Gel images were first quantified by SAFA (Das *et al*., 2005), and normalised by subtraction (equation 1). Upper and lower vigintiles were calculated from the base reactivities for all *furina* transcripts. Finally, reactivity 1 and 0 were defined by the upper and lower vigintiles for all base reactivities for a given RNA and values below zero were set to zero (equation 2). Shape reaction buffer: 333 mM HEPES, 3.3 mM Mg, 333 mM NaCl.

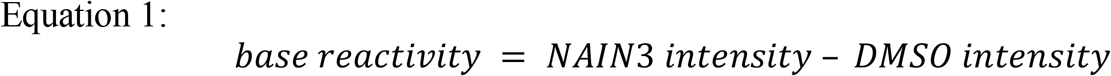

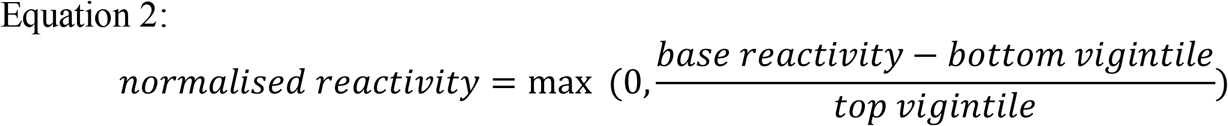

## Results

### Two furina transcript isoforms are expressed throughout embryonic development

Previous studies have demonstrated that FurinA proprotein convertase cleaves Nodal and facilitates establishment of left-right asymmetry (Constam and Robertson, 2000; Tessadori *et al*., 2015) Through analysis of zebrafish genomic databases, we find that in addition to the previously described *furina* transcript, a long *furina* variant X1 is also expressed during zebrafish embryogenesis. Variant X1 consists of an exceptionally large (∼4kb) 3’UTR sequence that has been only computationally predicted. Analysis of available RNA-seq datasets shows that during somitogenesis, novel *furina* variant X1 is expressed at levels higher than the previously identified short *furina* isoform ((White *et al*., 2017) and Figure 1A,B). To determine if variant X1 can be detected in embryos, we collected cDNA libraries from adult fish ovaries and 50% epiboly, 10 somite (10-som) and 18 somite (18-som) developmental stages and performed semi-quantitative PCR. In agreement with the RNA-seq data, we observed that the predicted variant X1 was not efficiently amplified in the ovary and at 50% epiboly. The highest expression levels are detected at 10-som and 18-som stages (Figure 1C). To quantitate the relative expression levels of the transcript isoforms, we performed a real-time PCR. We did not detect any significant difference between the expression levels of the two transcripts in the ovary, at the 1K cell stage or at 50% epiboly. Interestingly, there was significant upregulation of transcript X1 at 10-som, which reverts to similar levels by 18-som (Figure 1D). Next, we examined if there are differences in the expression pattern of the variant isoform in zebrafish embryos. Consistent with previous work (Tessadori *et al*., 2015) a *furina* coding region probe detected ubiquitous expression at all stages examined: 4-cell stage, bud, 10-som and 18-som. The expression pattern of variant X1 was similar to the Refseq-annotated transcript and there were no obvious differences observed. At 10-som and 18-som, we noticed that there were areas in embryo where both transcripts were slightly enriched. Thus, both *furina* transcripts are ubiquitously expressed, throughout embryonic development, with a slight enrichment in the eyes, hindbrain, and somite structures (Figure 1E).

**Figure 1.**
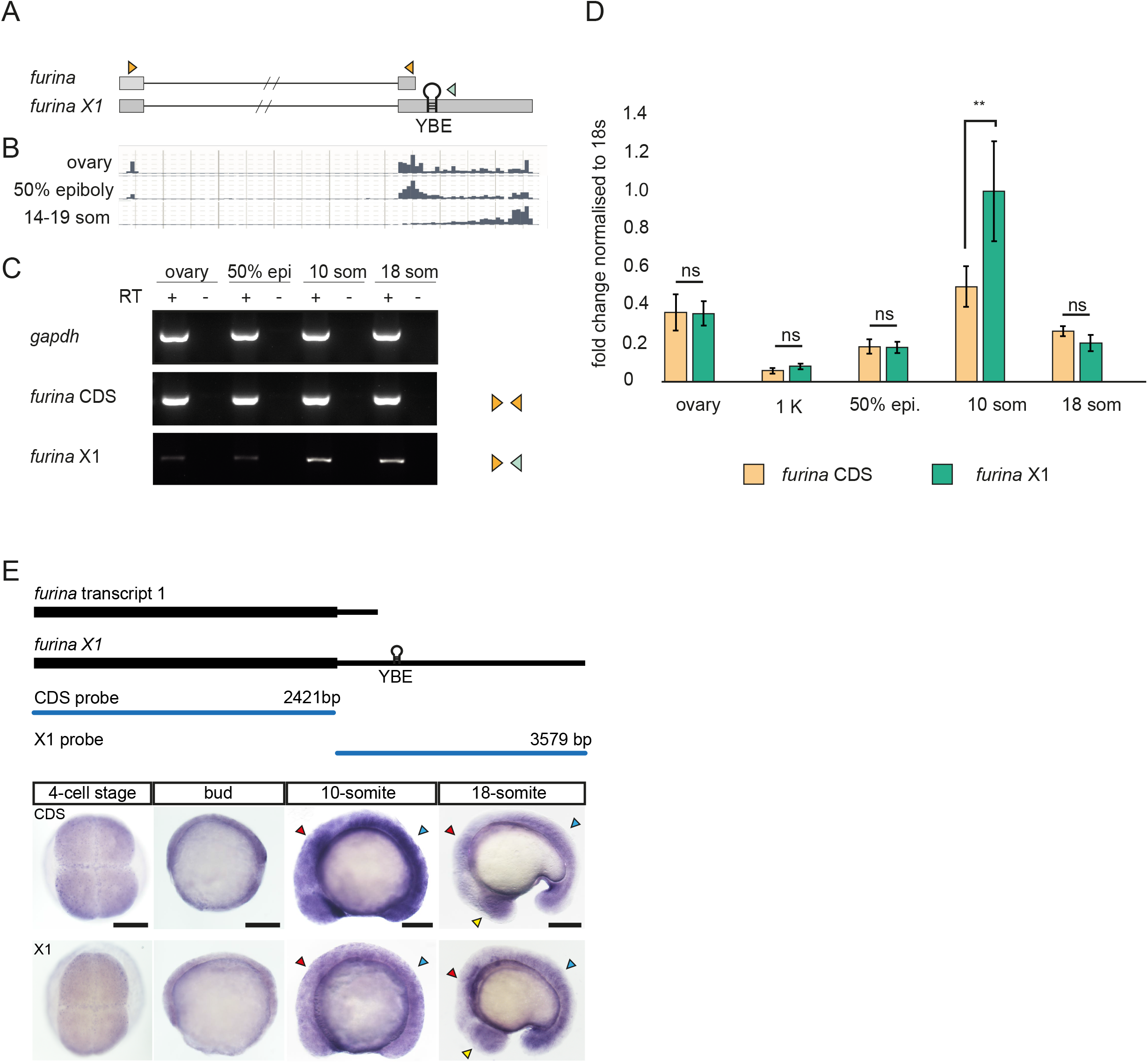
Zebrafish *furina* transcripts show different 3’UTR lengths, with a predominant long isoform expressed during somitogenesis. A) Schematic of *furina* transcript 1 and variant X1, orange arrowheads represent primers to amplify transcript 1, one orange and one green primer were used to detect variant X1. Hairpin indicates the YBE motif in the 3’UTR of X1. B)RNA-seq excerpt from ovary, 50% epiboly and 14-19 som stages. C)Semi-quantitative RT-PCR detecting *gapdh* positive control, *furina* coding sequences (CDS) and *furina* variant X1 3’UTR (X1). D)Quantitative PCR analysis from ovary, 1-K, 50% epiboly, 10 som and 18 som embryos. Orange bars represent *furina* coding region, green bars show exclusively variant X1. ** *p* < 0.01 in unpaired two-tailed student’s t-test. E)*Whole mount in situ* hybridization with distinct probes to detect transcript 1 or variant X1. Yellow arrows point to enrichment in eyes, red in hindbrain and blue in somites. Scale bars, 100 µm.

### The furina X1variant harbours the YBE element

We found that many nodal pathway components harbour a 3’UTR element containing an AGCAC sequence motif, followed by a stem and a variable length loop region (hairpin) (Zaucker *et al*., 2017). Computational analysis identified the *furina* variant X1 as a potential mRNA with a YBE element in the 3’UTR (Figure 2A). If the *furina* YBE motif behaves similarly to the 3’UTR motif in *lefty1*, they should co-localise in early zebrafish embryos. To test this hypothesis, *furina* variant X1 and *lefty1* fluorescent mRNA reporters were co-injected into 1-cell stage embryos and analysed for co-localisation at the 4-cell stage. The *furina* variant X1 fluorescent mRNA reporter co-localised with *lefty1* in 100% of injected embryos (n=11; Figure 2B). To determine if mutations affecting the 3’UTR YBE motif disrupted reporter mRNA localisation at the 4-cell stage, we generated deletion constructs that are predicted to change the YBE structure. Fluorescent reporter *furina* ∆ YBE had the entire YBE motif deleted within the 3’UTR. *Furina* Stem-break (SB) had a mutation disrupting base-pairing within the stem of the YBE and *furina* Stem-restore (SR) had a swapped sequence that restores base-pairing within the stem of the hairpin. Injected fluorescent LacZ-βglobin mRNA served as a negative control and sqt/ndr1 RNA as a positive control. Embryos were injected with a fluorescent reporter and then scored for reporter RNA localisation at the 4-cell stage. LacZ-βglobin fluorescent mRNA reporter distribution was diffuse everywhere in the blastoderm, but in contrast, *sqt/ndr1* RNA localised asymmetrically in one or two cells in 95% of embryos (n=19). *furina* variant X1 fluorescent reporter localised asymmetrically in 75% of embryos (n=17), whereas deletion of the YBE motif resulted in mis-localisation of the mRNA in 90% of embryos (n=18). Breaking of the stem caused 15% of SB-reporter injected embryos to show complete mis-localisation and 45% showed diffuse asymmetric distribution of the reporter. Injection with SR restored the localisation by 25% when compared to SB (Figure 2C). Thus, the furinaX1 variant harbours a functional YBE motif.

**Figure 2.**
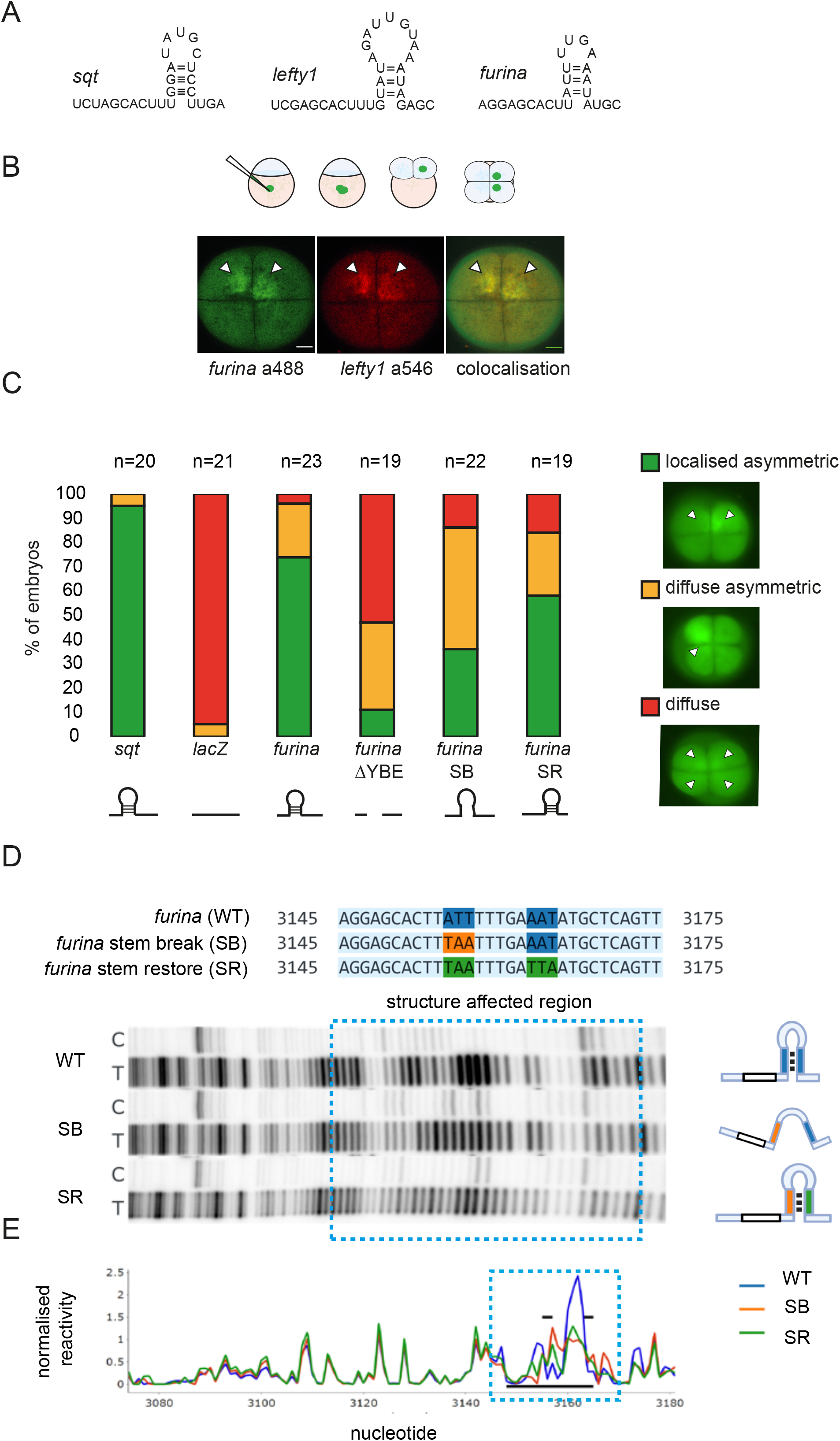
The *furina* variant X1 transcript harbours a YBE motif in the 3’UTR. A)Schematic showing hairpin YBE motifs in *sqt, lefty1* and *furina* variant X1. B)Top: schematic of localisation assay, embryos were injected in a yolk at the 1-cell stage and mRNA localisation was analysed at the 4-cell stage. Bottom: Confocal images showing co-localisation of *furina* (labelled with alexa 488-UTP) fluorescent reporter and *lefty1* (labelled with alexa 546-UTP) in zebrafish embryos at 4-cell stage. Scale bar, 20 µm. C)Stacked bar graph showing localisation categories in fluorescent reporter mRNA injected embryos: localised asymmetric (green), diffuse asymmetric (yellow), and diffuse (red). Schematic representations of fluorescent RNAs of control sqt, lacZ and furina variant X1 RNAs (full length furina or disrupting the YBE motif) are shown below in black. D)Structure probing of *furina* X1 3’UTR: structural mutations generated based on the predicted stem loop (SB, SR). Structure footprint gels for *furina* RNAs. Blue highlight sequences (*top*) and blue dotted box (bottom) indicates the structure affected region. Lanes labelled C indicate control untreated RNA that was reverse transcribed; lanes labelled T indicate RNA treated with NAI-N3 before reverse transcription. E)Quantification of structure footprint gels. Blue dotted box represents the structure affected region (blue highlighted sequences). Black bars indicate predicted YBE stem loop structure.

The *furina X1* UTR was also investigated by *in vitro* RNA structure-probing (Figure 2D,E). In *furina* WT, we observe a region of increased probe reactivity, flanked by regions of relatively low probe reactivity. Reactivity with the probe indicates high probability of single-strandedness, and this pattern approximately corresponds to the predicted stem-loop region in figure 2A. The pattern noticeably flattens in *furina* SB and is partially restored in *furina* SR (Figure 2D). Quantification of the gel footprints (Figure 2E) shows that RNA structure outside of the YBE stays largely unchanged in all RNAs tested, suggesting that structural mutations within the YBE only cause local changes in structure when probed *in vitro*. Comparing the quantified pattern of RNA structure by Pearson correlation (0.62 for WT vs SB and 0.84 for WT vs SR) indicates partial loss of the overall RNA structure in SB and partial restoration in SR. This partial loss and restoration of structure aligns with the *in vivo* reporter assays in figure 2C. Therefore, the furinaX1 variant harbours a functional YBE motif with a structured RNA motif.

### Ybx1 pull-down reveals interactions with the furina variant X1

The RNA-binding protein Ybx1 binds to the YBE motif in the 3’UTR of nodal and lefty mRNAs and forms a YBE/Ybx1 translational repression and RNA localisation module (Kumari *et al*., 2013; Zaucker *et al*., 2017). We hypothesised that the YBE we identified in the furina X1 variant RNA is likely to bind to Ybx1. To test if Ybx1 interacts with *furina* X1 mRNA *in vivo*, we performed RNA-immunoprecipitation with zebrafish embryonic lysates from various stages (50% epiboly, 10-som and 18-som) using anti-Ybx1 antibodies, followed by RT-PCR (Figure 3A,B). In western blots, we detect Ybx1 protein at 50% epiboly, 10-som, and to lesser extent at 18-som. Pull-downs with mouse IgG served as a negative control (Figure 3B). Consistent with our published work, *lefty1* was enriched at 50% epiboly and 10-som stage. Using primers to distinguish between the *furina* coding sequence versus 3’UTR sequences, we find that at 50% epiboly and 10-som there is enrichment of *furina* long isoform over mouse IgG controls in pull-downs in both set of samples, whereas no significant furina is detected in pull-downs from 18-som embryo lysates (Fig 3C). Ribosomal 5S amplification served as a negative control in all cases, and *lefty1* served as positive control. These results indicate that furina mRNA is in a complex with Ybx1 protein at 50% epiboly and at 10-som, but not at 18-som.

**Figure 3.**
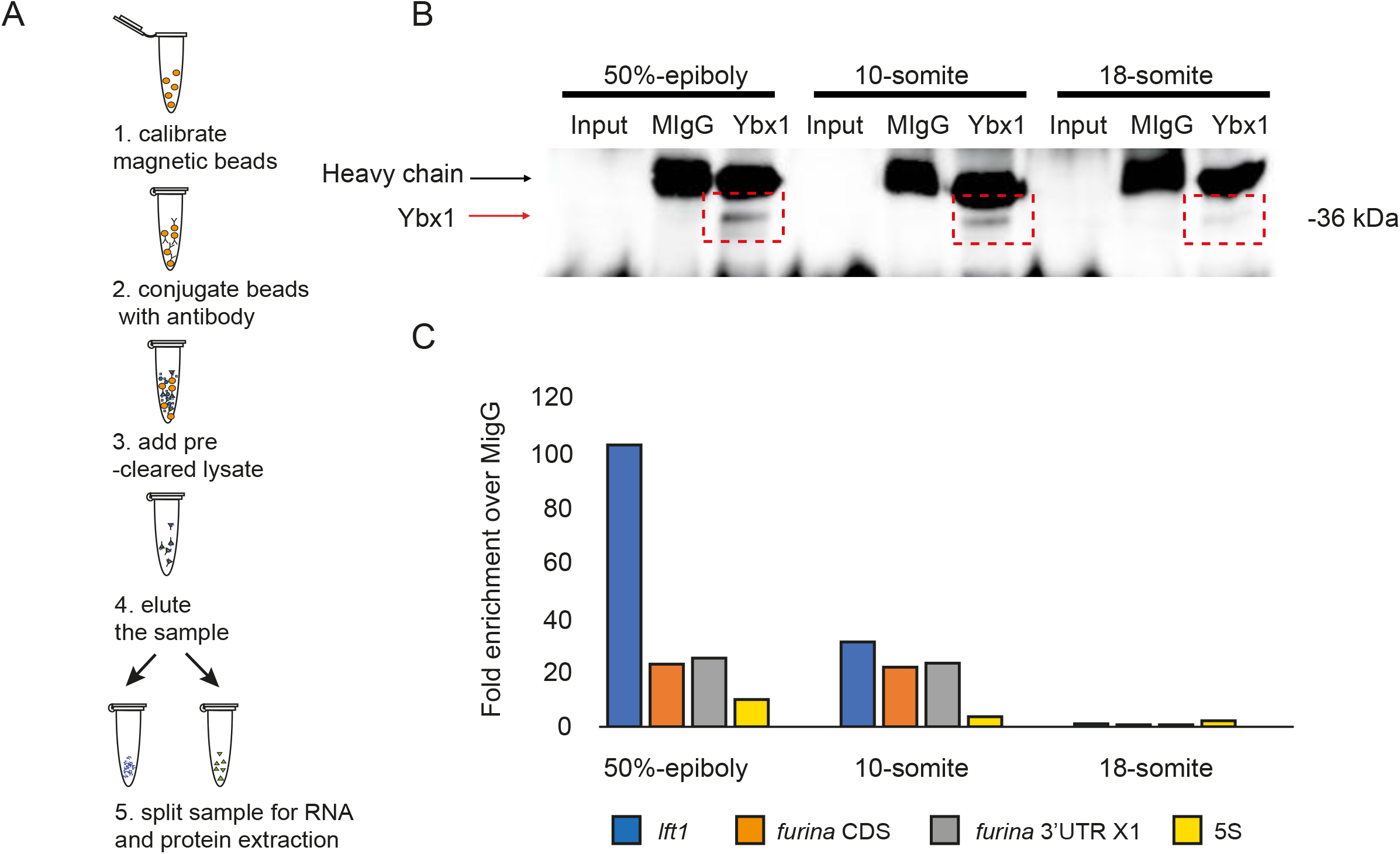
Ybx1 protein interacts with *furina* variant X1 mRNA. A)Schematic of RNA-IP of embryo lysates with anti-Ybx1 antibodies. B)Representative Western Blot showing pull down of Ybx1 from 50% epiboly, 10-som and 18-som stage embryonic lysates; black arrow indicates IgG heavy chain; red arrow indicates Ybx1 band. C)qPCR analysis of samples from IP; Fold enrichment normalised to input and mIgG control are shown for l*efty1* (blue), *furina* coding region (orange), *furina* variant X1 3’UTR (grey) and 5S negative control (yellow).

### Variant X1 of *furina* is translationally repressed by Ybx1 in zebrafish embryos

The YBE in *nodal* and lefty1/2 functions in translational control of these RNAs during early development. To test if the *furina* variant X1 is similarly regulated by Ybx1, we performed our established translation assay that utilises *ybx1*^sa42^ temperature sensitive mutant fish to measure the expression of microinjected reporter constructs in *ybx1* mutants by circumventing the early embryonic death phenotype of ybx1 null mutants (Kumari *et al*., 2013). We injected *ybx1* mutant embryos at 1 cell stage with *furina-*sfGFP:*furina3’UTR* RNA, Ybx1 protein function was disabled as shown in the schematic and embryos were analysed for protein expression (Figure 4 A and B). Analysis of the *furina*-GFP reporter showed a strong band in maternal (M)*ybx1* mutants, but not in control embryos or un-injected controls. We quantified the expression levels based on band intensity and normalised to Actin levels. Significantly higher FurinA GFP levels were detected in lysates from *ybx1* mutant embryos (Figure 4C,D).

**Figure 4.**
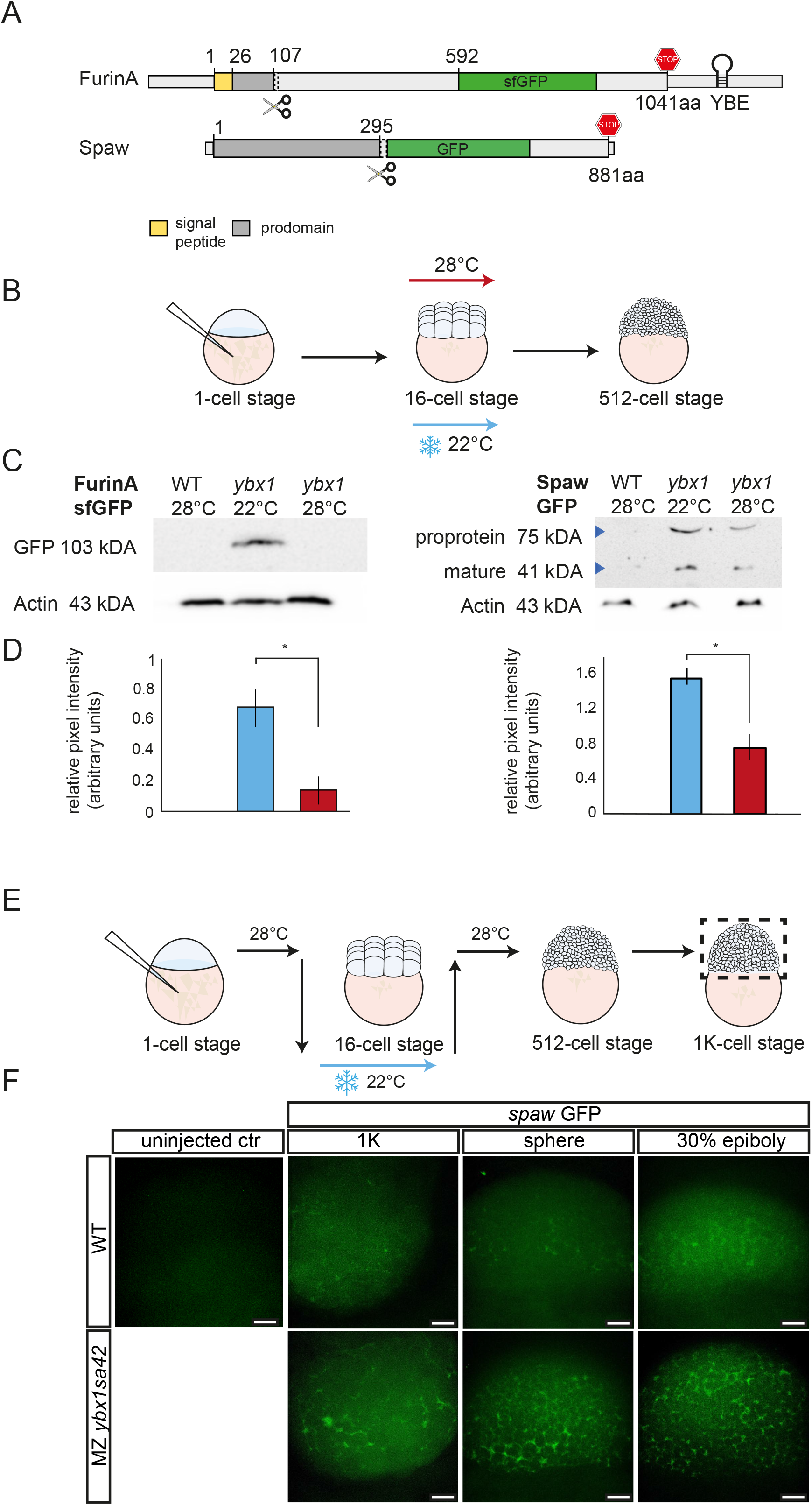
FurinA translation and Spaw maturation is increased in *ybx1* mutant embryos. A)Schematic of GFP fusion reporters for *furina* X1 and *spaw*. Hairpin indicates the YBE motif in furina RNA. The FurinA pro-domain is shown in dark grey and the signal peptide in yellow; for Spaw, the pro-domain is indicated in dark grey; stop codon indicated in red in furin and spaw. B)Schematic of temperature shift of *ybx1*^*sa42*^ mutant embryos. Embryos were injected with GFP reporter-fusion mRNA, grown at 28 ºC until the 16-cell stage, shifted to 22 ºC until the 512-cell stage, when protein was extracted. C)Western blots on embryonic lysates to detect showing FurinA-sfGFP (left) and Spaw-GFP fusion protein (right), following injection of embryos with mRNA reporter. Lanes were loaded with lystaes from un-injected wild type control, M*ybx1* at 22ºC and M*ybx1* control embryos at 28ºC. Actin loading control is aligned to the corresponding samples. D)Bar graphs show quantification of corresponding band intensities, with FurinA-sfGFP (left) and mature Spaw (right) normalised to the Actin loading control, *p* *<0.05. E)Schematic showing temperature shift of *ybx1* mutant embryos following injection with *spaw* GFP reporter. F)Confocal images of 1-K, sphere and 30% epiboly wild-type (top panel) and *ybx1* mutant embryos (bottom panel). Scale bar, 50 µm.

As FurinA cleaves Spaw/Nodal to control Nodal signalling range in the lateral plate mesoderm (Tessadori *et al*., 2015), we posited that overexpression of FurinA in *ybx1* mutant embryos would lead to increased processing of Spaw and aberrant Nodal signalling in *ybx1* mutant embryos. To test this, we injected *spaw*-GFP fusion mRNA into *ybx1* mutants at the one-cell stage, followed by temperature shift at the 16-cell stage, and collected embryonic lysates at 500-cell stage (Figure 4 A,B). We detect two Spaw protein products, consistent with Spaw precursor (75 kDa) and a mature Spaw (41 kDa) (Figure 4C). Interestingly, in mutant lysates, we found elevated levels of Spaw mature peptide compared to controls. Quantification of band intensity of Spaw mature domain (n=3 experiments) and normalisation to Actin loading control (Figure 4D) show that in the absence of Ybx1 function, FurinA protein levels are upregulated. Therefore, Ybx1 translationally represses *furina* variant X1 mRNA.

Mature Nodal/Southpaw is secreted into the extra-cellular space. We therefore examined Spaw maturation in *ybx1* mutant embryos over time (Figure 4E,F). We injected *spaw* GFP reporter at the 1-cell stage and analysed expression at the 1K, sphere and 30% epiboly. At the 1K, sphere and 30% epiboly stages, both intra and extra cellular GFP expression was detected by imaging the blastoderm of wild type embryos (Figure 4E,F). By contrast, in *ybx1* mutant embryos, there was an obvious increase in accumulation of Spaw-GFP in the extracellular space (Figure 4F). We measured total Spaw-GFP fluorescence levels across the blastoderm in wild type and *ybx1* mutant embryos and found that overall fluorescence levels were similar (Supplementary Figure S1). Therefore, even though overall Spaw-GFP levels are apparently similar in both wild type and mutant embryonic blastoderms, Spaw maturation is increased and faster in *ybx1* mutants, resulting in excess and premature secreted mature Nodal protein in mutant embryos.

### Mutant ybx1^sa42^ embryos exhibit left-right asymmetry defects

Increased processing of Spaw/Nodal and altered Nodal levels are associated with left-right (LR) asymmetry defects (Long et al., 2003; Tessadori et al., 2015). Therefore, we next investigated if *ybx1* mutant zebrafish embryos show defects in LR asymmetry. WISH at 18-som shows a 15% increase in a bilateral or right sided spaw expression in *ybx1* mutants (Figure 5A,B, n=44). Abnormal *spaw* expression is associated with organ laterality defects, such as defective looping of the heart and mis-positioning of the liver, pancreas, and gut (Long, Ahmad and Rebagliati, 2003). In *ybx1* mutant embryos, we detected a 20% increase in an aberrant looping of the heart compared to both wild-type embryos at 22°C, and 28°C *ybx1* controls (Figure 5C,D). A proportion of the temperature-shifted *ybx1* mutant embryos also showed no looping, or complete inversion of the heart chambers (Figure 5C,D). Analysis of heart looping in *ybx1* mutant embryos showed altered heart morphology in mutant embryos compared to wild type controls. Further examination of *bmp4* and *notch1b* expression in the heart showed that in addition to the looping defects, *ybx1* mutants also have a larger heart (n=8/24) and AV canal (n=4/29), respectively (Figure S2).

**Figure 5.**
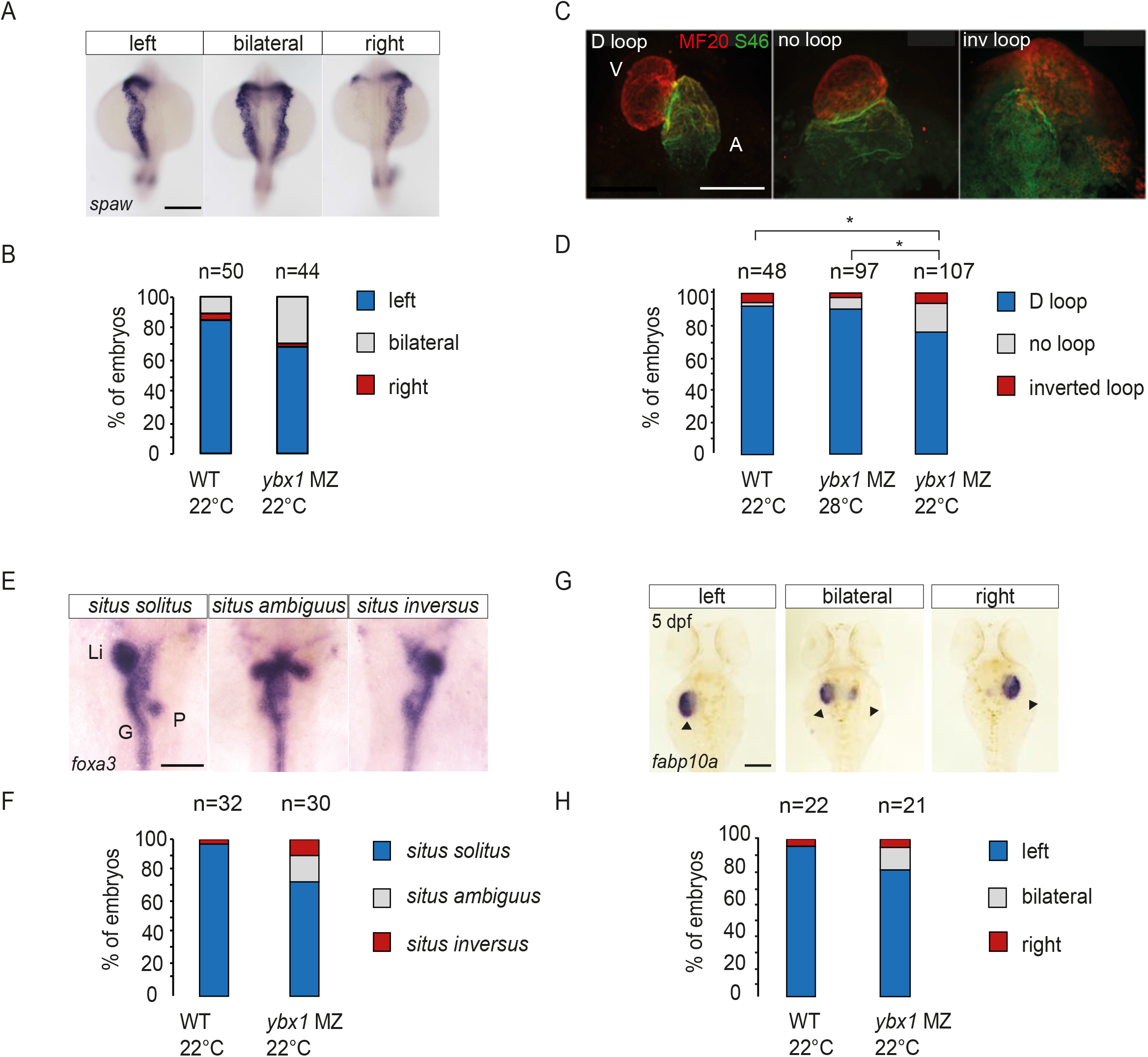
Mutant *ybx1*^*sa42*^ embryos have abnormal left-right expression. A)WISH to detect *spaw*/*ndr3* in wild type and *ybx1*^*sa42*^ mutant embryos. B)Bar graph shows proportion of embryos with spaw expression on the left (blue), bilateral (grey) or right (red). Scale bar, 100 µm. C)Immunofluorescence showing MF20 labelling in the ventricle (red) and S46 labelling in the atrium (green). Representative D loop, no loop and inverted (inv) loop hearts are shown; Scale bar, 100 µm. D)Quantification in graph shows looping of heart: Directional loop (D loop, blue), no loop (grey) and inverted looping (red) of the heart, with *ybx1* mutant embryos at 22°C showing higher % of inverted and no loop hearts compared to control WT or *ybx1* embryos at 28°C. *p* *<0.05. E)WISH to detect *foxa3* at 55 hpf in wild type and *ybx1* mutant embryos, Li indicates liver, P is pancreas and G is gut. Scale bar, 100 µm. F)Quantification of visceral organ positioning in wild type and *ybx1* mutant embryos shows increase in *situs ambiguous* (grey) and *situs inversus* (red) with decrease in *situs solitus* (blue) in *ybx1* mutant embryos. G)Expression of *fabp10a at* 5 dpf, arrowheads indicate liver. Scale bar, 100 µm. H)Quantification of liver positioning in wild type and *ybx1* mutant embryos showing *situs solitus* (blue), *situs ambiguous* (grey) and *situs inversus* (red).

To assess the positioning of visceral organs such as liver, gut and pancreas, we examined expression of *foxa3*. In *ybx1* mutants, we observed increase in either duplication, mispositioning or inversion of the liver and pancreas (Figure 5 E,F). By contrast, the majority of control embryos showed correct organ positioning (Figure 5E,F). Similarly, *fabp10a* expression revealed a similar trend where the majority of control embryos had a liver on the left-hand side and 20% of the *ybx1* mutant embryos had either duplicated livers or inverted liver position (black arrowheads, Figure 5G,H). Therefore, left-right organ positioning is affected in *ybx1* mutant embryos.

### Cardiac morphogenesis is affected in ybx1^sa42^ mutant embryos

To determine if heart morphogenesis is altered in the *ybx1* mutant embryos, we performed immunofluorescence with the ZN8 antibody, which labels the AV canal, and quantified the AV canal width. At 55 hpf, we observed an overall increase in width of the AV canal compared to WT embryos. Additionally, wild type controls had more consistent measurements, whereas *ybx1* mutant embryos presented a broad range in AV canal width (Figure 6A-C). This raised the possibility that *ybx1* mutant embryos may harbour defects in heart valves and cardiac blood flow.

**Figure 6.**
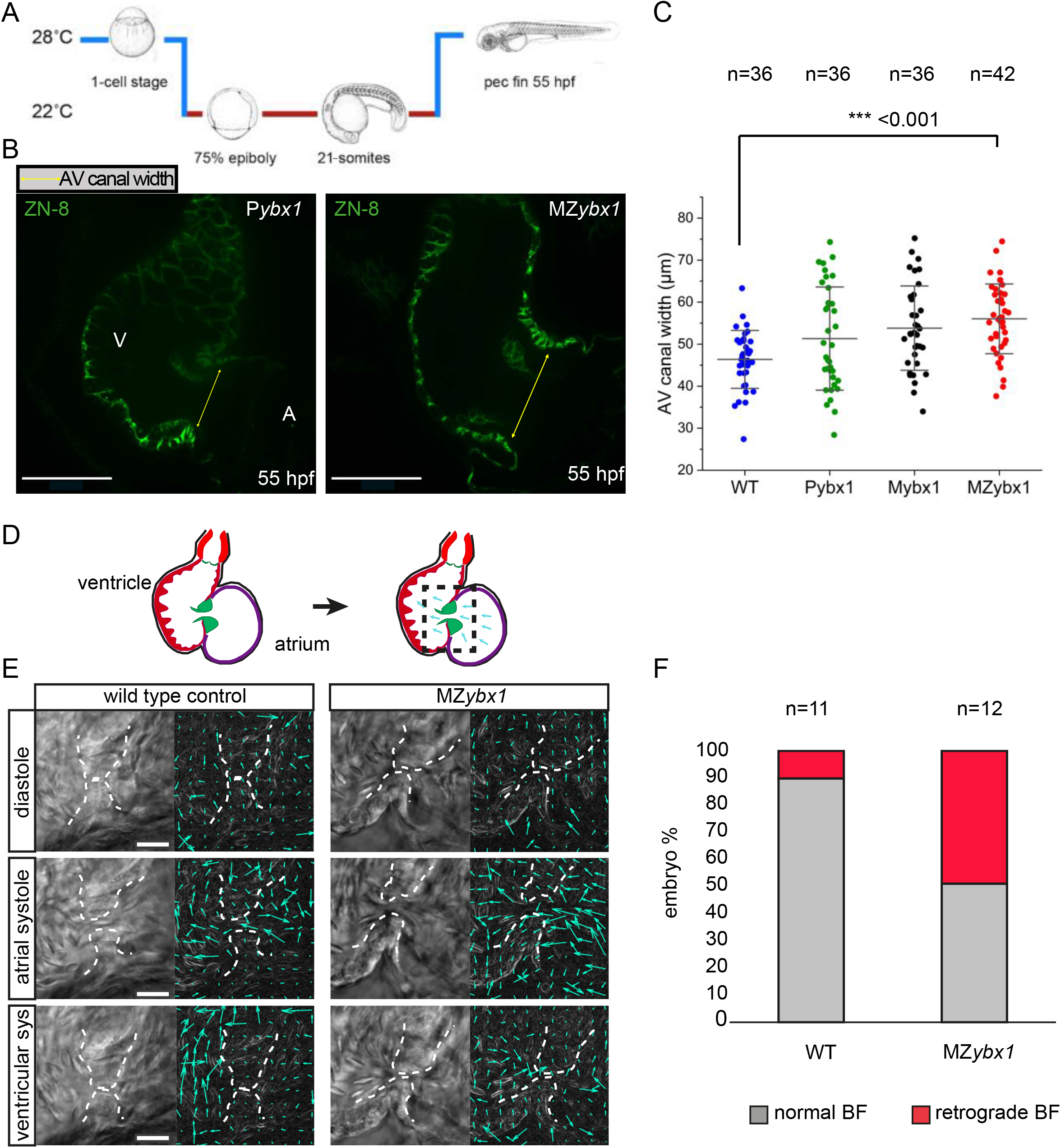
*Ybx1* mutant embryos show abnormal heart morphogenesis. A)Schematic of temperature shift; *ybx1* mutant embryos and controls were shifted to 22 ºC at 75% epiboly until 21-som and then grown at 28 ºC until 55 hpf. B)Confocal images of zebrafish heart showing immunofluorescence with the ZN-8 antibody to label the of atrioventricular (AV) canal. Yellow arrows indicate the width of the AV canal in control P*ybx1* and MZ*ybx1* mutant embryos. V, ventricle and A, atrium. Scale bar, 25 µm. C)Quantification AV canal width in µm from immunostaining of wild type (blue), P*ybx1 (*green), M*ybx1* (black) and MZ*ybx1* embryos (red), *p* ***<0.01. D)Schematic of zebrafish heart at 5 dpf (left) and schematic of PIV analysis (right); black dotted box shows area of heart imaged and analysed. E)Left panels, DIC images from movie of wild type and *ybx1* mutant hearts during diastole, atrial systole, and ventricular systole. On the right PIV analysis of the images are shown, cyan arrows indicate movements of red blood cells; Scale bar, 20 µm. F)Percent embryos showing normal blood flow (BF) versus retrograde BF in wild type and *ybx1* mutant embryo groups.

To determine the consequences of the larger AV canal in *ybx1* mutants, we analysed the heartbeat of wild type and *ybx1* mutant embryos by live-imaging at 5 dpf (Figure 6D). The movement of red blood cells was tracked, and the direction of blood flow during the contraction of the heart muscle was assessed. Live-imaging at different stages of the heartbeat cycle and tracking of the red blood cells at 5 dpf revealed that 6 out of 12 *ybx1* mutant embryos show red blood cells going back to the atrium, in contrast to wild type controls where blood flows from the atrium to the ventricle (Figure 6E,F and and Supplementary Movies 1,2). Quantitation showed that over 50% of *ybx1* mutants show retrograde blood flow (Figure 6F), which is consistent with abnormal heart morphogenesis in these embryos.

### Mutant *ybx1* embryos manifest heart valve regurgitation and reversed blood flow

Analysis of the *ybx1* heartbeat revealed that during the ventricular systole, instead of blood cells being pushed out of the ventricle, a large number of red blood cells leak through the heart valves and move back into the atrium. To examine blood flow patterns in these embryos in-depth, we analysed and quantified the movements of the red blood cells around the atrioventricular canal (Figure 7A). We focused on two timepoints during ventricular contraction in *ybx1* embryos with reverse blood flow (RF) versus wild type controls. At the beginning of the ventricular systole, red blood cells move in the same direction in both wild type and mutant embryos (Figure 7 B,D). However, at the end of the ventricular systole, in mutant embryos we observe that red blood cells change direction and start moving back, whereas in wild type embryos the movement trajectory remains the same (Figure 7 C,E). Analysis of the velocity of the red blood cells during contraction of the ventricle confirmed that in *ybx1* RF embryos there is increase in cell movement back into the atrium, when in control embryos little movement is detected in this direction (Figure 7F). Furthermore, in *ybx1* embryos, initially the cells move out of the ventricle, but over time their velocity gradually decreases. In control embryos, however, movement of cells out of the ventricle remains steady and consistent (Figure 7G). The altered movement of blood cells is consistent with our observations that *ybx1* mutants have a larger AV canal (Figure 6B,C), heart valves appear misaligned, and result in reverse blood flow, showing a heart defect at 5 dpf. This shows the important role of translational control by Ybx1 protein, not only in the left-right asymmetry establishment during early embryogenesis in zebrafish, but also in the correct heart morphogenesis and heart valve function.

**Figure 7.**
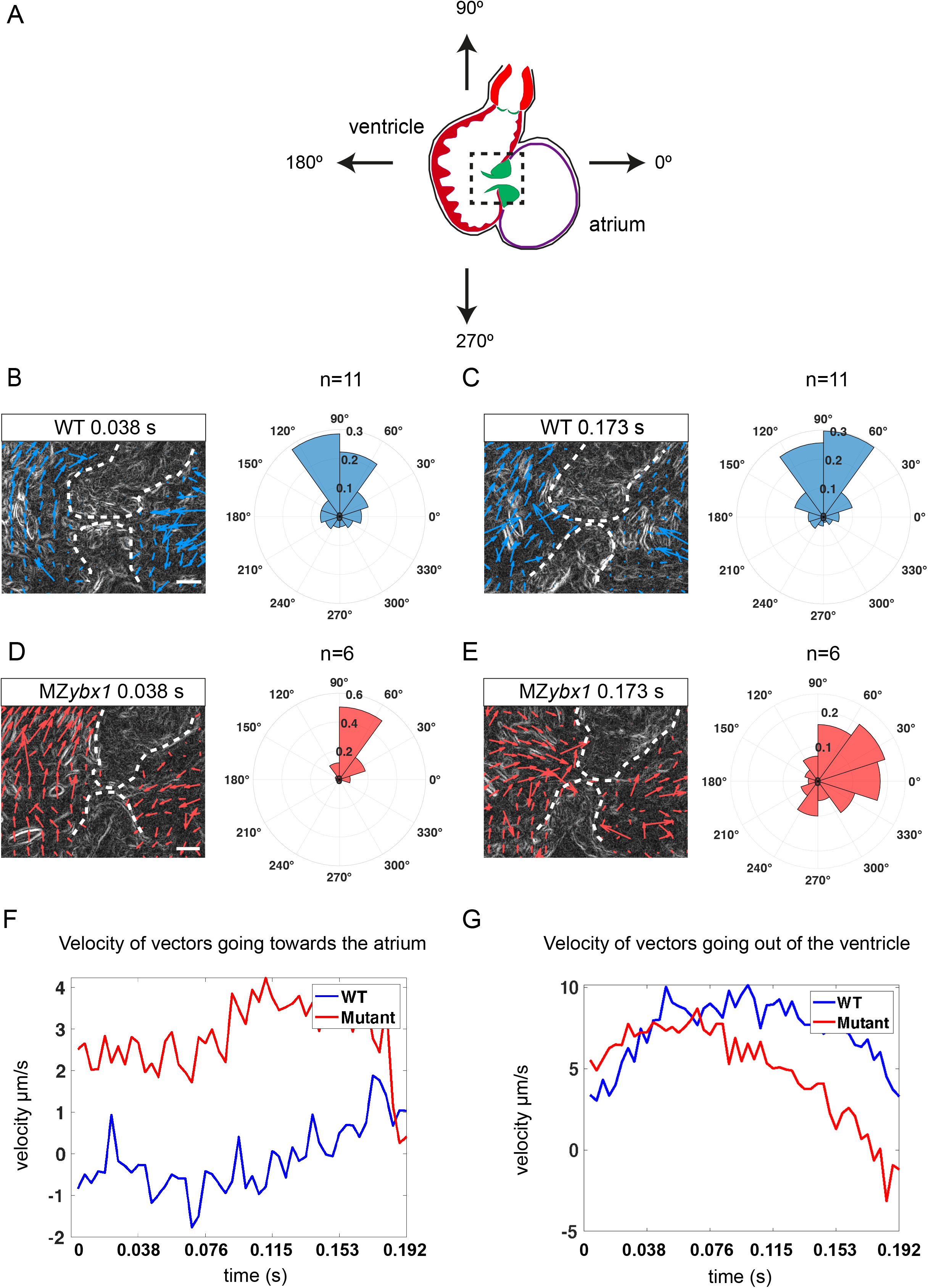
Mutant *ybx1* embryos have retrograde blood flow at 5 dpf. A)Schematic of zebrafish heart at 5 dpf; Arrows indicate direction of flow; Black box shows region analysed in PIV. B)Representative image of a wild type embryo, showing PIV application at 0.038 s of ventricular systole. Vectors generated from red blood cells are shown as blue arrows. Rose plot shows quantification of direction of the blood flow at 0.038 s for the imaged wild-type hearts (n=11). Scale bar, 25µm. C)A PIV image of wild-type ventricular contraction at 0.173s (left) and quantification of direction of generated vectors (rose plot on right) (n=11). Vectors are shown as blue arrows. Scale bar, 25µm. D)A PIV image showing the direction of blood flow in MZ*ybx1* embryos with retrograde blood flow at 0.038s. Vectors are shown as red arrows. Rose plot shows analysis of direction of blood flow in MZ*ybx1* embryos with retrograde blood flow (n=6). Scale bar, 25 µm. E)PIV analysis at 0.173 s of ventricular systole and quantification of vector direction in *ybx1* embryos with retrograde blood flow at 0.173 s (n=6). Vectors are shown as red arrows. Scale bar, 25 µm. F)Analysis of average velocity of horizontal vectors moving towards the atrium over the duration of ventricular contraction. Wild type embryos are shown in blue and *ybx1* mutants in red. Units, µm/s. G)Analysis of average velocity of vertical vectors moving out of the ventricle during ventricular contraction. Wild type embryos, blue and *ybx1* mutants, red. Units, µm/s.

## Discussion

In this work, we describe a novel *furina* variant X1 mRNA with a YBE element in the 3’UTR. Similar to other *nodal* mRNAs, we show that *furina* YBE motif is recognised by Ybx1 RBP and has an important role in translational repression of FurinA. Previous work by other groups showed that FurinA protein cleaves Spaw and establishes its signalling range (Tessadori *et al*., 2015). We observed that in the absence of Ybx1, FurinA is premature and overexpressed and, as a result, Nodal/Spaw processing is increased. This causes excess Spaw protein levels and ectopic Spaw expression outside the left lateral plate mesoderm. This leads to left-right asymmetry defects, with abnormal heart looping and abnormal positioning of liver and pancreas. Our work thus identifies a novel mechanism of regulation of left-right asymmetry and cardiac morphogenesis through the RNA-binding protein Ybx1. Although many transcription factors have been found to control left-right patterning across animal species, very few RNA-binding proteins and/or corresponding RNA elements have been identified in this process (Rago *et al*., 2019; Vejnar *et al*., 2019), and therefore, our work can lead to potentially new clinical diagnostic tests for LR asymmetry defects that are currently unassigned.

RNA elements have a crucial regulatory role in mRNA biogenesis, stability, localisation and transation. The Hu RBP family in human neurons is a great example. The HuR, HuB, HuC and HuD proteins preferentially bind to ‘AU’ and ‘U’ rich sequences in 5’ and 3’UTRs of the RNAs. These interactions are key in regulating RNA stability and translation in changing cellular conditions (Lopez de Silanes *et al*., 2004). In early zebrafish embryos, 3’UTR elements and translational control play crucial roles in regulation and stability of maternal RNAs (Mishima and Tomari, 2016; Winata *et al*., 2021). Mammalian studies also identified general motifs via CLIP-seq that some RBPs bind to, for instance: UGAU, AAUAAA. Interestingly, the IGF2BP3 family of RBPs recognises CWWCATCA and TGCACTAT, to facilitate mRNA transport and enhance stability (Li *et al*., 2017). In this study, we experimentally validated the YBE motif in *furina* mRNA by *in vitro* RNA structure probing. Although we do not see a perfect recapitulation of the hypothesised 3 bp stem loop structure, we see some support for RNA structure in that region and note that our *in vitro* observations might be confounded by the lack of stabilising trans-acting factors that would normally be present *in vivo*. One possibility is that perhaps there is RNA structure heterogeneity in the region, where all these RNAs can generate a structure similar enough to that observed by structure probing, but with WT *furina* having the highest propensity and SB *furina* the lowest propensity for the predicted structure.

We show that Ybx1 interacts with *furina* mRNA at the embryonic stages prior to organ morphogenesis. The phenotype observed in *ybx1* mutant embryos is likely due to a lack of translational control of *furina* mRNA and is consistent with overexpression of FurinA protein. When there is an excess of FurinA protein, cleavage of Spaw is upregulated and the midline of an embryo is saturated with Spaw, which causes bilateral expression and consequently left-right asymmetry defects (Tessadori *et al*., 2015). To date, there are very few studies showing involvement of RNA-binding proteins in left-right asymmetry. One study described translational repression of Bicc1 on *dand5* 5’UTR mRNA and showed a role for this interaction in the establishment of laterality (Maerker *et al*., 2021). Dand5 is a negative inhibitor of Nodal signalling and probably acts at a different level of the Nodal pathway, downstream of *furina* and Ybx1 regulation. Interestingly, we observed the left-right asymmetry defects in approximately 20% of the mutant embryos. Ybx1 is a global translational repressor and it also interacts with the Nodal inhibitors Lefty1 and Lefty2 (Zaucker *et al*., 2017; Sun *et al*., 2018). Although in the *ybx1* temperature sensitive allele the mutation was triggered in late epiboly stage, there is still upregulation of Lefty1 in *ybx1* mutants. During somitogenesis Lefty1 is expressed at the midline of the embryo preventing Spaw to cross the midline barrier and ensuring its leftward expression (Lenhart *et al*., 2011). However, in *ybx1* mutants, upregulation of Lefty1 activity likely represses elevated Spaw. This, as a result, might reduce the severity of left-right asymmetry phenotype in *ybx1*^*sa42*^ embryos.

In addition to the left-right organ positioning defects, we also found that the AV canal in the *ybx1* mutants is enlarged and consequently the embryos have retrograde blood flow. This phenotype could also potentially be independent of or in addition to the interaction of FurinA with Spaw. FurinA is a ubiquitously expressed protein that cleaves a range of targets, which are required for correct heart and AV canal development, such as Notch1B in zebrafish (Zhou *et al*., 2021). In *Xenopus* and mouse it was shown to interact with the Bone morphogenic pathway (BMP) family, also important for heart development (Cui *et al*., 1998; Constam and Robertson, 2000). The lack of translational control of *furina* in *ybx1* mutants might also disrupt interactions with one of these targets and lead to additional heart defects. It is also possible that Ybx1 might regulate heart morphogenesis through other Nodal-independent mechanisms (Noël *et al*., 2013).

There are reports of regulation of the heart development by non-coding molecules such as microRNAs (miRNAs). Key cardiac development genes such as *tbx2, notch1* and *cspg2* are regulated by miRNAs in zebrafish (Kalayinia *et al*., 2021). Studies in mice have shown that increased levels of miR-25 leads to depressed cardiac function, but inhibition of miR-25 reversed the phenotype and increased survival in a mouse heart failure model (Wahlquist *et al*., 2014). Studies from patients have shown that reduced amounts of miR-26a, miR-30b and miR-195 in stenotic valves led to increased chance of developing valve leaflet fusion. These micro RNAs are thought to be involved in modulating calcification genes and when their levels are lower, calcium begins to accumulate at the aortic valve leaflet and changes the morphology (Nigam *et al*., 2010).

We find that *ybx1* mutant embryos have retrograde blood flow at 5 dpf. Retrograde blood flow can be observed normally up to day four in larvae, and is important for modelling of the AV canal and valve development, and mutations in the heart valve precursor gene *klf2a* cause aberrant valve development (Vermot *et al*., 2009). In *ybx1* mutants, we observe that red blood cells squeeze through the valves back to the atrium. This is similar to a human condition called mitral valve regurgitation, which is characterised by a leaky valve leading to backflow of blood. The extent of the symptoms varies, with some patients being asymptomatic and others experiencing dizziness, breathlessness, and irregular heartbeat (Adams, Rosenhek and Falk, 2010). The expansion of the atrioventricular canal likely prevents heart valves from shutting properly, and during ventricular contraction the blood cells squeeze between the valves and go back to the atrium of mutant hearts. Therefore, our *ybx1* mutant can serve as a zebrafish model for this disorder and could potentially lead to better understanding and future treatments of heart valve regurgitation disorders.

## Acknowledgements

We thank members of the KS lab for feedback and comments on the manuscript, the Warwick BSU aquatics facility staff for fish care, and Balasubramanian laboratory for support and advice.

## Supplemental Figure Legends

**Supplementary figure S1.**
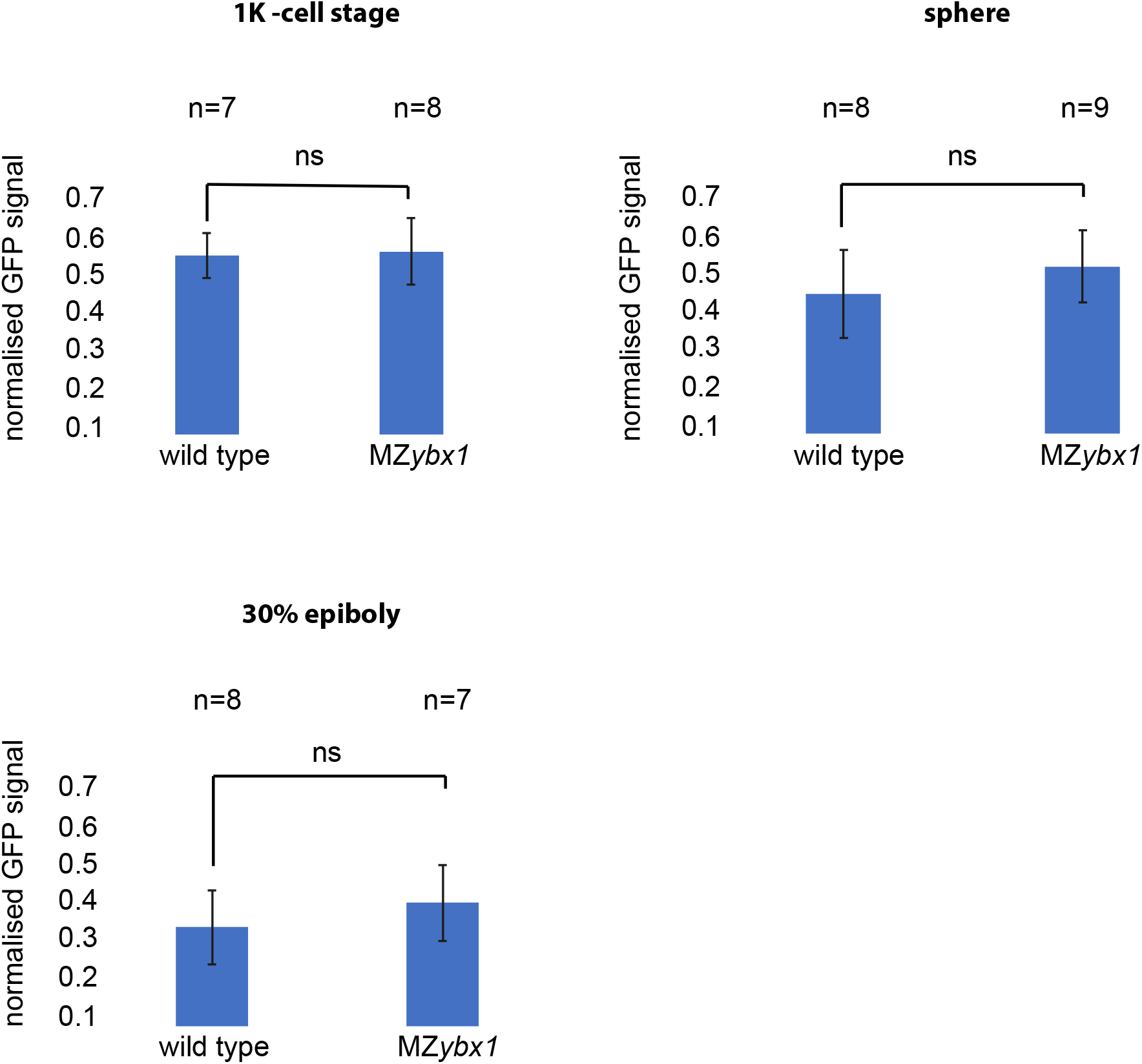
Total Spaw-GFP fusion protein fluorescence levels are similar in wild type and *ybx1* mutant embryos. Quantification of fluorescence levels across the entire blastoderm at 1K, sphere and 30% epiboly; Spaw-GFP signal was normalised to mCherry mRNA reporter control.

**Supplementary figure S2.**
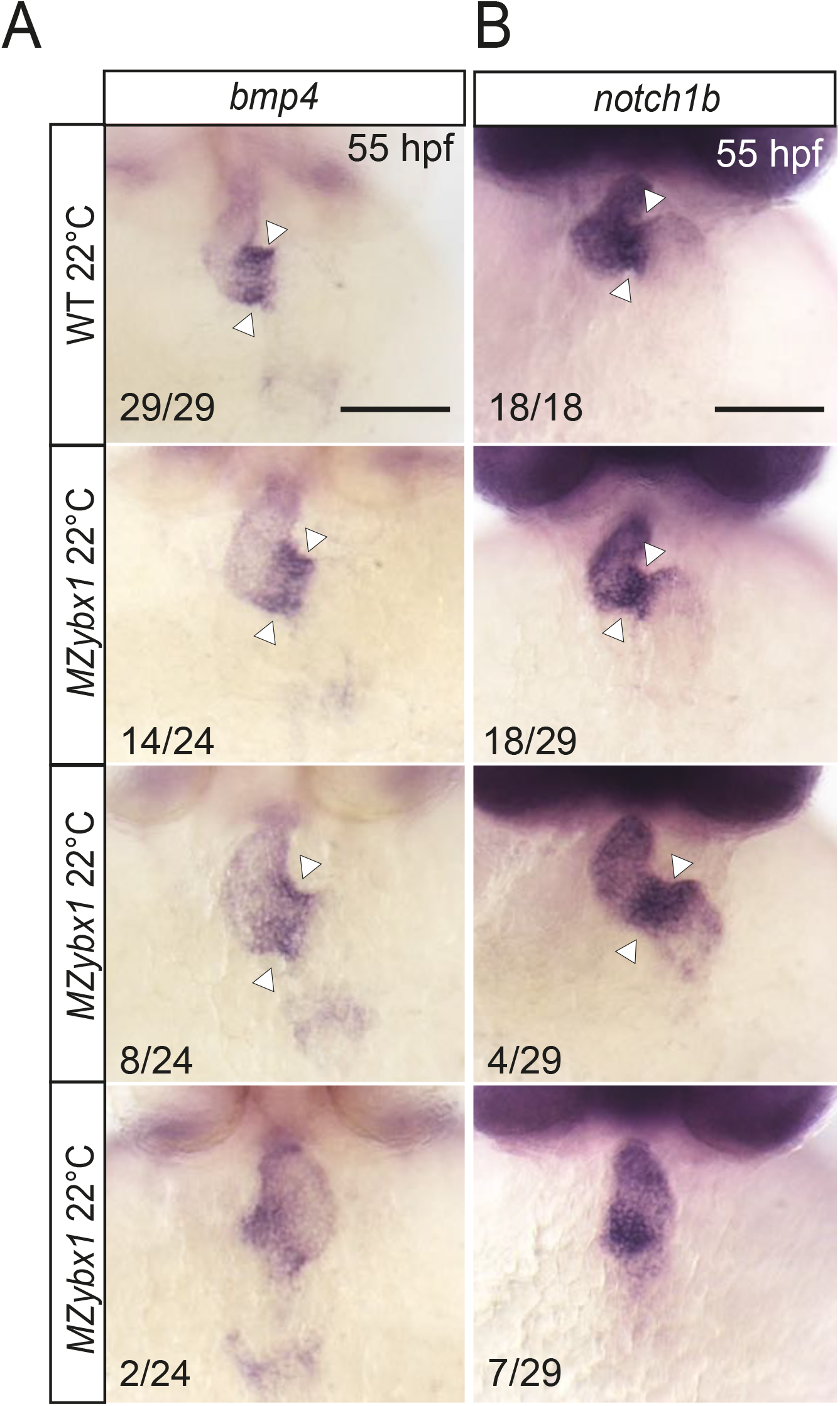
The heart appears enlarged in MZ*ybx1* mutant embryos at 5 dpf. A)WISH showing *bmp4* expression in wild type and *MZybx1* mutant embryos. White A)arrowheads point to the AV canal. Scale bar, 100 µm. B)WISH with *notch1b* probe, showing expression in the AV canal at 55 hpf in wild type and *MZybx1* mutant embryos. White arrowheads point to the AV canal, scale bar, 100 µm.

## Supplemental Movie legends and Graphical abstract

**Supplementary Movie 1**

DIC movie showing heartbeats in a 5dpf wild type zebrafish embryo (left) and PIV analysis (right) showing direction of blood flow; Cyan arrows indicate red blood cells; Images were captured at 13 frames per second intervals, V marks ventricle and A marks atrium; Scale bar 20 µm.

**Supplementary Movie 2**

DIC movie of heartbeats in a 5 dpf MZ*ybx1*^*sa42*^ mutant embryo showing retrograde blood flow (left), and PIV analysis (right) showing the movement of the red blood cells. Images were captured at 13 frames per second intervals, V marks ventricle and A shows atrium. Scale bar 20 µm

## Graphical abstract

Schematic of model showing Ybx1 translational regulation of *furina*, facilitating left-sided *nodal* expression during somitogenesis. In the absence of Ybx1 binding to *furina* mRNA, there is translational de-repression leading to increase in FurinA protein levels, premature and increased Nodal protein maturation which then diffuses across the midline and leads to bilateral *nodal* transcript expression.

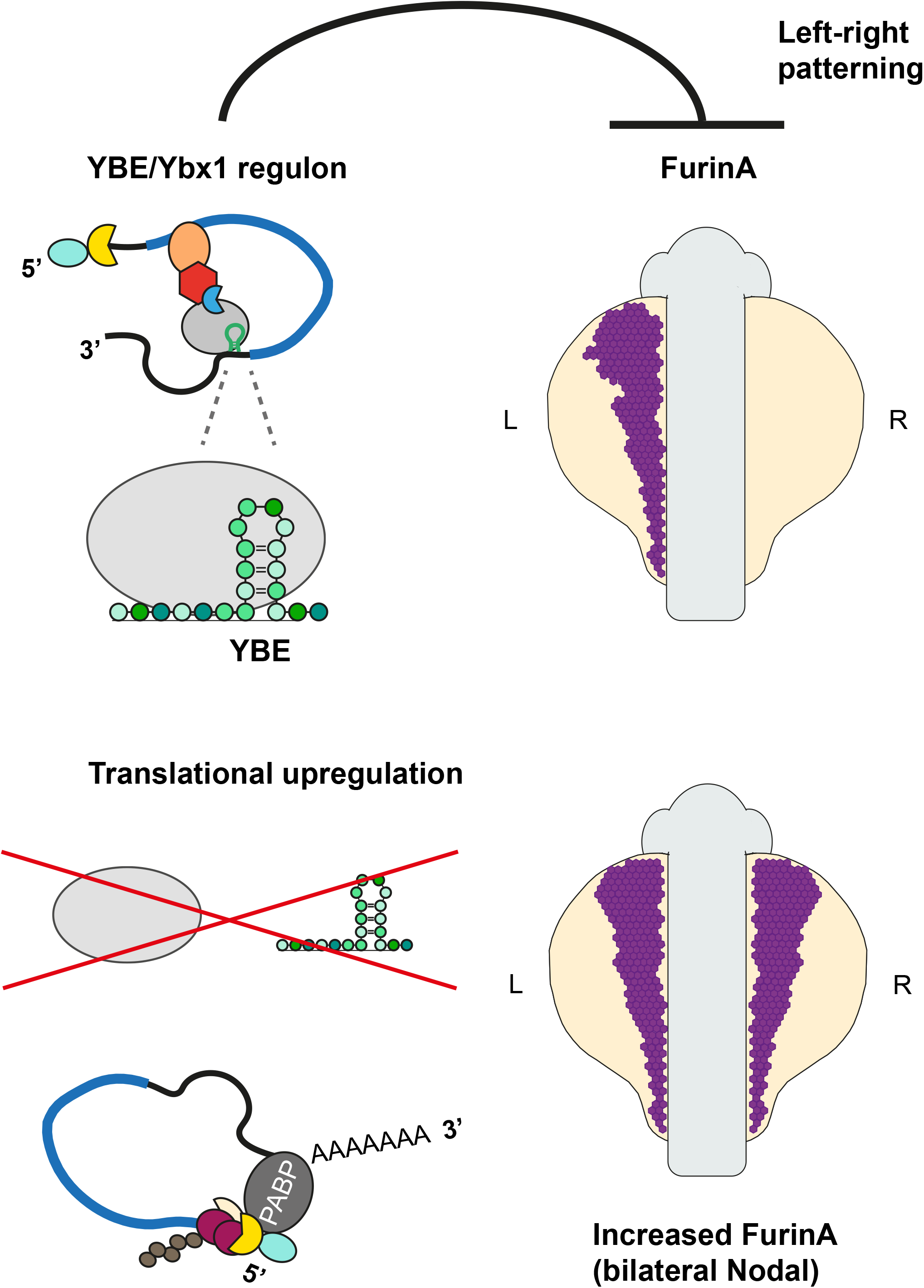

